# The SARS-CoV-2 Spike harbours a lipid binding pocket which modulates stability of the prefusion trimer

**DOI:** 10.1101/2020.08.13.249177

**Authors:** Loic Carrique, Helen ME Duyvesteyn, Tomas Malinauskas, Yuguang Zhao, Jingshan Ren, Daming Zhou, Thomas S Walter, Julika Radecke, Jiandong Huo, Reinis R Ruza, Pranav NM Shah, Elizabeth E Fry, David I Stuart

## Abstract

Large trimeric Spikes decorate SARS-CoV-2 and bind host cells via receptor binding domains (RBDs). We report a conformation in which the trimer is ‘locked’ into a compact well-ordered form. This differs from previous structures where the RBD can flip up to recognise the receptor. In the ‘locked’ form regions associated with fusion transitions are stabilised and the RBD harbours curved lipids. The acyl chains bind a hydrophobic pocket in one RBD whilst the polar headgroups attach to an adjacent RBD of the trimer. By functional analogy with enteroviral pocket factors loss of the lipid would destabilise the ‘locked’ form facilitating receptor attachment, conversion to the postfusion state and virus infection. The nature of lipids available at the site of infection might affect the antigenicity/pathogenicity of released virus. These results reveal a potentially druggable pocket and suggest that the natural prefusion state occludes neutralising RBD epitopes, achieving conformational shielding from antibodies.

**Highlights:**

- **SARS-CoV-2 Spike can adopt a ‘locked’ conformation with all receptor binding domains (RBDs) down, likely to represent the prefusion resting state**
- **This ‘locked’ conformation is compact and stable, braced by lipid bound within a potentially druggable pocket**
- **Key neutralization epitopes are shielded in the ‘locked’ form**
- **Loss of lipid may trigger a cascade of events that lead to cell entry analogous to the role of lipids in enterovirus cell entry**

## Introduction

Severe acute respiratory syndrome coronavirus-2 (SARS-CoV-2), the causative pathogen of COVID-19, emerged in late 2019 and has caused unprecedented global damage to human health and the economy (Nicola et al., 2020). Coronaviruses (CoVs) are pleomorphic enveloped RNA viruses possessing an unusually large positive sense, single stranded genome. They are classified into four genera, the α-, β-, γ- and δ-coronaviruses (Fehr and Perlman, 2015). SARS-CoV-2 belongs to group β, alongside two other viruses responsible for recent disease outbreaks, MERS-CoV (2012) and SARS-CoV-1 (2001-2003) (Gorbalenya et al., 2020). Although many coronaviruses are not pathogenic to humans, zoonotic coronaviruses have repeatedly jumped species barriers. Furthermore coronaviruses are prone to recombination, making the search for cross-protective antivirals and vaccines urgent (Su et al., 2016).

The SARS-CoV-2 viral envelope is decorated with glycosylated Spike, a large (>1200 residues per protomer) trimeric type I fusion protein. The majority of neutralising antibodies appear to target Spike and it is the primary focus of attention in the development of non-classical (e.g. RNA, DNA and VLP based) vaccines (Amanat and Krammer, 2020). The protein has at least two major functions. The first is to attach to receptor(s), with ACE2 identified as having high affinity for one domain of Spike, termed the receptor binding domain (RBD), and many neutralising antibodies function by abrogating this interaction (Wu et al., 2020). The second function is to act as the machine which fuses the viral and host cell membranes allowing the viral genome to enter the cell (Rey and Lok, 2018). This multistage process is not fully understood but it is thought that receptor attachment destabilizes the prefusion state and cleavage of Spike by host protease into two major fragments, S1 and S2, releases the metastable protein to undergo massive conformational changes, discarding the S1 portion which includes the RBD. This allows S2 to convert into an extended helical bundle projecting a fusion peptide outwards to engage the host cell membrane before then hauling the membranes together to merge cytoplasmic and viral compartments. The use of enzymatic cleavage to release the potential energy of metastable fusion machines is common to many enveloped viruses, and coronaviruses belong to a very large set, bearing so-called ‘type I fusion proteins’, which share a common helical bundle mechanism, but diverge widely in cell attachment methods (Rey and Lok, 2018). Structural studies to date have largely used a highly engineered version of Spike (benefitting from experience with stabilising SARS-CoV-1 and MERS Spikes) to obtain a stable prefusion form, which although non-functional, maintains the authentic antigenicity (Pallesen et al., 2017). The most pertinent change was the introduction of two proline residues (2P) in the hairpin region linking the HR1 and Central helices (Figure S1) that become a single helix in the postfusion state. The prolines prevent formation of the extended helix and were characterized as stabilizing the prefusion Spike (Pallesen et al., 2017). This engineered ‘prefusion’ Spike has a rather loose structure, with significant portions of the trimer subunit disordered, and is found with the RBDs (residues 333-527) arrayed in different configurations of two orientations, either ‘up’ or ‘down’ (PDBIDs:6VSB, 6VYB, 6VXX, 6X2A,B,C,9) (Henderson et al., 2020; Walls et al., 2020; Wrapp et al., 2020). The RBD sits top-centre on the Spike, surrounded by the larger NTD, also part of S1. In the ‘up’ conformation the ACE2 binding site is accessible, however this conformation is thought to destabilize the Spike, facilitating fusion with the host cell membrane (Yan et al., 2018). It remains a puzzle how the Spike interrogates its surroundings for receptors whilst maintaining integrity of the prefusion state, however the accepted model has been that RBDs stochastically flip up to transiently reveal the ACE2 binding site (Yuan et al., 2017). This view of the Spike structure has been challenged by the observation that purified wild-type Spike ectodomain exists as a mixture of just two states: a ‘locked’ (all RBDs down) prefusion state not previously seen and a postfusion conformation (Cai et al., 2020).

Efforts to meet the need for COVID-19 small molecule therapeutics have been primarily directed against essential viral proteases and polymerases (Khan et al., 2020; Zhang et al., 2020), with much interest, and some success, centred on opportunities for drug repurposing (Yin et al., 2020). An alternative approach is antibody therapy, either through the use of convalescent serum or monoclonal antibodies (Duan et al., 2020; Huo et al., 2020). By analogy with the search for AIDS therapeutics we would expect that, if there is sufficient will and resource, effective antivirals will be found that target a range of proteins, including non-enzyme targets, as exemplified by, for HIV, several clinically approved entry inhibitors (Esté and Telenti, 2007). Also repurposed drugs have been found that neutralize Ebolavirus by destabilising the functional equivalent of Spike (Zhao et al., 2016) and for coronaviruses EK1, a mimetic of part of the S2 fragment of Spike, has been shown to act as an entry inhibitor (Xia et al., 2019). Understanding Spike structure and conformational changes is therefore important to define the anatomy of the target not only for neutralizing antibodies (Jiang et al., 2020), but also new or repurposed chemical entities (Fleishman et al., 2011).

Here, we report a high-resolution cryo-EM structure for the 2P version of SARS-CoV-Spike ectodomain, revealing a tightly packed arrangement of the Spike trimer with all RBDs down. This ‘locked’ form orders substantial portions of structure including regions adjacent to the fusion peptide, suggesting it hinders conversion to the postfusion state. Remarkably, the ‘locked’ conformation, which is probably dis-favoured by the proline mutations, is stabilized by hydrophobic ligands, probably one or a mixture of several unsaturated fatty acids, buried in a hydrophobic pocket in the RBD. Stabilisation is achieved by RBD-RBD cross-linking by the head group of the lipid and also by contacts mediated by conserved glycans. Consistent with this observation, we show that lipid-like molecules modulate the stability of the Spike. These data reveal that the RBD of SARS-CoV-2 Spike has a potentially druggable pocket and suggest alternative models for Spike dynamics and receptor attachment from those currently accepted.

## Results

### Locked prefusion SARS-CoV-2 Spike dis-favours conversion to postfusion state

A detailed analysis was performed of cryo-EM data collected during a study of the binding of the neutralizing antibody CR3022 (Huo et al., 2020) to engineered Spike ectodomain, which has residues K986 and V987 replaced with prolines (2P), an abrogated furin cleavage motif and a C-terminal foldon domain to maintain trimeric association (Wrapp et al., 2020) (Methods, Figures S1-S3, Table S1). This revealed an further Spike conformation (referred to here as ‘locked’) in addition to the ‘up’ or ‘down’ configurations previously observed (predominantly two, one with all three RBDs down (‘all down’) and the other with one RBD rotated up and two down (‘one up’) (Walls et al., 2020; Wrapp et al., 2020) (Methods, Figure S2). The ‘locked’ configuration, comprising <10% of the total picked particles, constitutes the most tightly packed form observed for this SARS-CoV-2 Spike protein (key features are shown in Figure 1A-E). This is reflected in the increased number of residues (a total of 1,089 for the 1247 residue ectodomain construct) that can be fitted into the density map, especially in the NTD. Compared to the ‘looser’ all-down version of the trimer (PDB: 6VXX (Walls et al., 2020)) an extra 120 residues are visualized and the additional stability of the protein has permitted a high-resolution analysis (3.1 Å) from just 36,000 particles. In both the ‘all-down’ and the ‘locked’ conformations the RBDs point downward in a 3-fold symmetric arrangement. Whilst they are broadly similar in structure, there are significant differences. The comparison shows that for the ‘locked’ form, the NTDs are rotated by approximately 7° and slide inwards towards the RBDs by ~ 7-10 Å. The RBDs in turn are shifted towards the central axis by 2-3 Å (the distance between the N-termini of adjacent RBDs decreases by almost 6 Å) (Figure 1A, Video S1). The tighter packing around the 3-fold axis locks the NTD loops into a more ordered structure, as reflected in the improved density maps. RBD packing around the 3-fold axis also leads to the ordering of a segment of S2 (T827-F855) (Figure S4) where Y837 interacts with the main chain of the adjacent protomer residues P589’ and R634’ (’ denotes neighbouring protomer) and the side chains L841 and F855 form a hydrophobic patch with residues V551’, T553’ and F592’. This region of S2 is adjacent to the fusion peptide and we suggest that this structure in the ‘locked’ form helps secure the prefusion state of the molecule.

**Figure 1.**
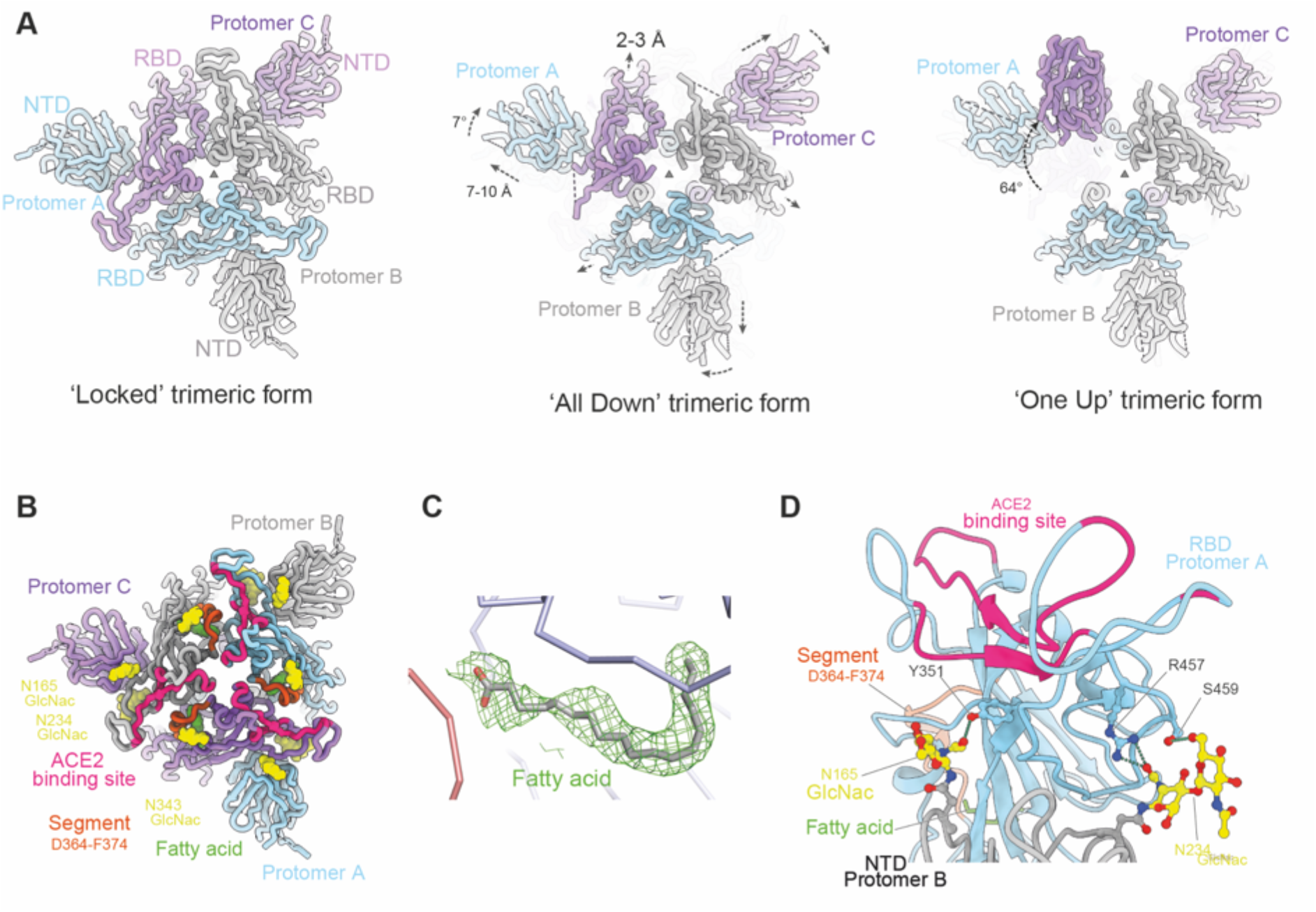
Overall structure of locked trimeric structure of SARS-CoV-2 Spike. (A) Comparison of the lipid bound Spike with the previously observed RBD ‘all down’ (PDB ID, 6VXX) and ‘one up’ Spike structures (PDB ID, 6Z97). The protomers are coloured in pale blue, grey and purple respectively. Relative rotation and translation of the NTD and RBD are indicated. (B) Top view of the RBD and NTD domains of the ‘locked’ Spike showing the relative positions of the bound fatty acids (green surfaces), the ACE2 binding sites are shown in red, α2 helices that have largest movement in brown, glycans at N165 and N234 of the NTD and N343 of RBD as yellow surfaces. (C) Density of the bound fatty acid. (D) Closeup of the interactions between glycans (yellow) on N165 and N234 of protomer B NTD (grey) and protomer A RBD (cyan).

In the ‘locked’ conformation, the ‘2P’ mutations K986P and V987P, introduced to prevent conversion to the postfusion conformation (Pallesen et al., 2017), are close to residue D427 (Figure S4). D427 is part of a conserved patch on the RBD that ‘caps’ the 986-987 hinge region, preventing premature conversion to the postfusion form (Zhou et al., 2020). In the wild-type Spike, K986 could form a salt bridge with D427’ to stabilize the ‘locked’ form but these salt bridges would be lost in the more open forms of the Spike (Figure S4). This latch in the wild-type Spike suggests that the ‘locked’ conformation is a more realistic representation of the ‘resting’ prefusion Spike conformation present at the viral surface, with the previously reported ‘all-down’ and ‘one-up’ structures transient conformations occurring just prior to ACE2 engagement, perhaps following a cellular cue. Consistent with this, a recent structural analysis of wild-type Spike lacking the 2P mutation revealed a similar such ‘locked’ state, with no sign of the more open prefusion Spike structures reported for the 2P stabilized molecule (Cai et al., 2020), however the coordinates are not available precluding a detailed comparison.

Glycosylation is often considered to mask potentially immunogenic protein epitopes. Three conserved glycosylation sites, located at N165, N234 and N343, have been reported for stabilized SARS-CoV-2 Spike expressed in 293F cells (Watanabe et al., 2020). All three lie close to RBDs in the ‘locked’ conformation, well positioned to shield the ACE2 binding site from immune recognition. In addition, the first or second glycosidic rings of the N165 and N234 glycans make polar interactions underneath the tip of the RBD, opposite to the ACE2 binding site: GlcNacN_165_ with Y351’ and GlcNac_N234_ with R457’ and S459’ (Casalino et al., 2020) (Figure 1D). We would expect these to be recapitulated in the highly processed sugars of wild-type virus (Watanabe et al., 2020).

### The ‘locked’ conformation is stabilized by a pocket factor which cross-links the RBDs

Careful inspection of the ‘locked’ structure revealed the presence of elongated continuous density extending deep inside the hydrophobic core of the RBD (Figures 1 and 2). The density occupies a curved pocket lodged between the ß-sheet and outer α-helices of some 589 Å^3^ volume (Figure 2) not found in the previously reported ‘all-down’ and ‘one-up’ structures. The length of the density is consistent with an 18-carbon fatty acid such as stearic acid, and its curvature suggests an omega-3 or omega-6 fatty acid with at least two unsaturated carbons (Figure 1C). In the absence of a formal chemical analysis of the molecule we have modelled it as stearic acid, however note, by analogy with enterovirus pocket factors, that there might be a mixture of lipids (Smyth et al., 2003). The fatty acid fits almost entirely within the pocket (85% of the surface area buried) and is stabilized by a series of hydrophobic interactions (Figure 2E-F), whilst the polar/charged headgroup is exposed at the end of the pocket where it faces a neighbouring RBD, interacting with R408’ and indirectly with Q409’ (Figure 2F). Interestingly, while the tip of the RBD, which includes the ACE2 binding site, is highly variable across β-coronaviruses, the residues composing the hydrophobic pocket are largely conserved (Figure 3). Of the 27 residues shielding the lipid, 23 are conserved and for none of the variable residues does the side-chain interact directly with the lipid. This strong conservation suggests that the pocket could be a common feature across β-coronaviruses. Residues D364-F374 move outwards, from the positions seen in previous structures, by ~6 Å to form the pocket entrance (Figure 2A-B, Video S2) adopting a conformation that has never previously been observed for the engineered 2P prefusion Spike (Walls et al., 2020; Wrapp et al., 2020) or the isolated RBD domain (for instance PDBIDs: 6VXX, 6VSB, 6VYB) (Figure 2C), but may be consistent with the form reported recently for the wild-type prefusion Spike (Cai et al., 2020) (coordinates not available). This reorganization allows favourable packing against the neighbouring RBD, especially *via* Y365 and Y369.

**Figure 2.**
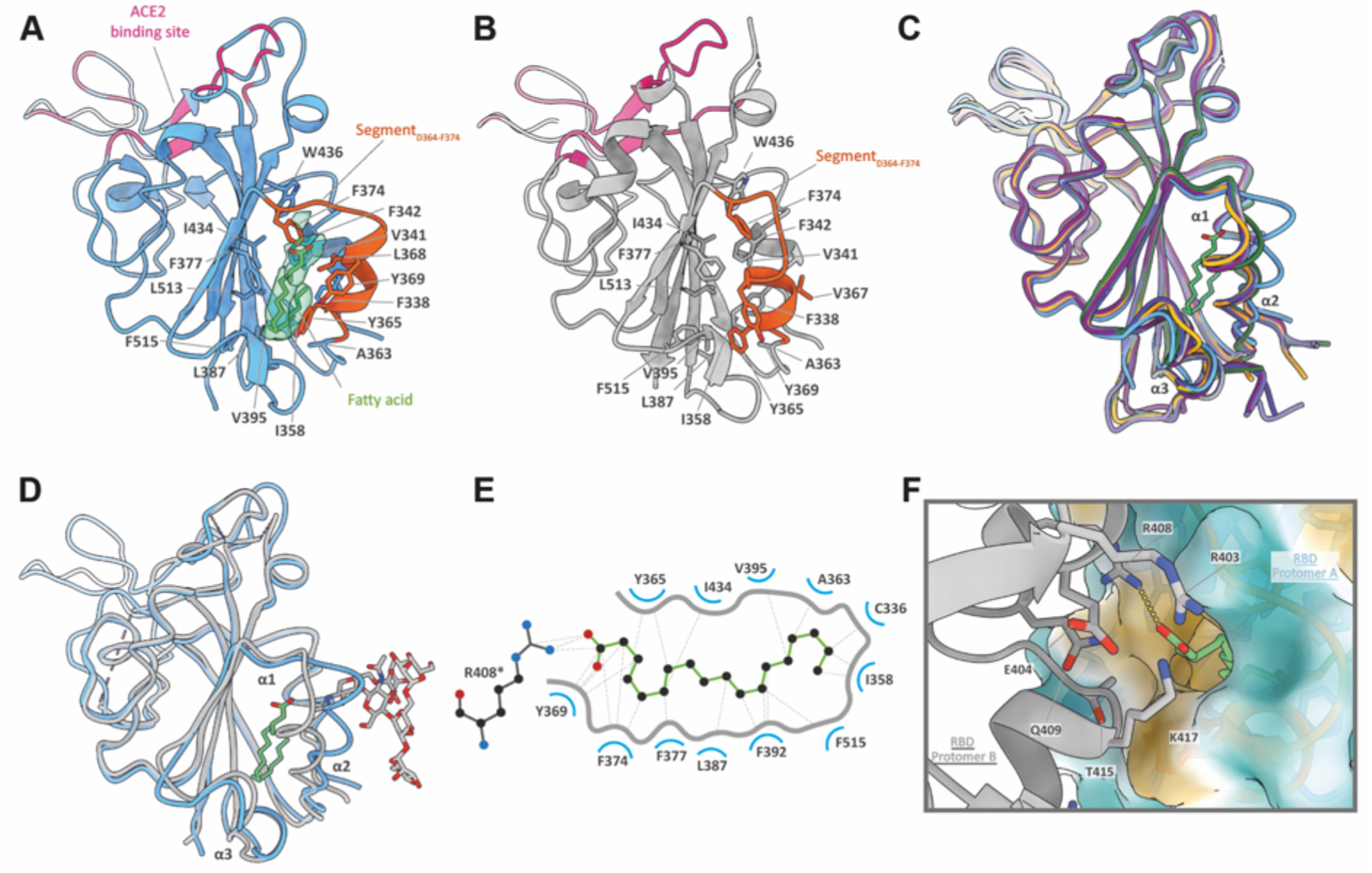
Details of the fatty acid binding site. (A) The fatty acid binding site in the locked Spike RBD. The polypeptide chain is drawn as blue ribbons, ACE2 epitope in red, and the segment with large conformational difference compared with the known Spike structures in orange. The fatty acid is shown as green sticks with density as surface, and protein side chains in contacts with the lipid as blue and orange sticks. (B) The collapsed fatty acid binding pocket of the RBD in the ‘all-down’ form of Spike. (C) Comparison of the fatty acid (green sticks) bound RBD of the ‘locked’ Spike (blue) with six crystal structures of RBD complexed with either ACE2 or Fab or nanobody (PDB IDs, 6YLA, 6ZCZ, 6M0J, 6LZG, 6YZ5 and 7BZ5). (D) Overlay of the fatty acid bound RBD (blue) with that from S309 Fab bound Spike (grey; PDB ID, 6WPS) showing the clashes of α2 helix with the glycans (grey sticks) that are part of the S309 binding epitope. (E) Ligplot showing the interactions of the lipid with residues lining the pocket. Hydrophobic interactions and hydrogen bond are shown as dotted lines. Residue R408 from a neighbouring RBD marked with a *. (F) The head group of the bound fatty acid is stabilized by polar or charged residues from neighbouring RBD.

**Figure 3.**
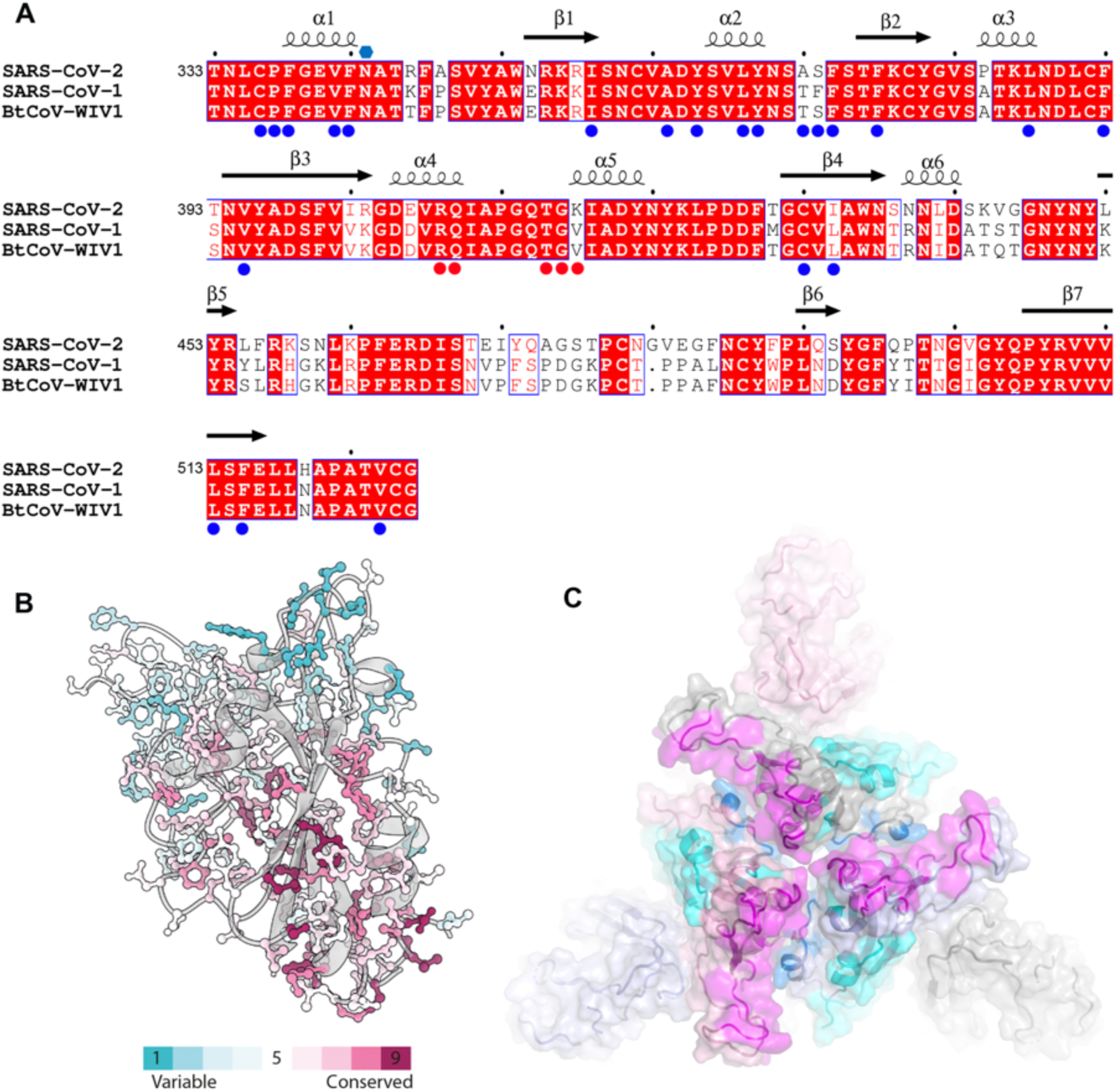
Conservation of the lipid binding site in the beta linage of coronaviruses and the hidden RBD epitopes of the locked Spike. (A) Sequence alignment of RBDs of three beta-coronaviruses. Conserved amino acids are shown in red background, residues shielding the fatty acid marked by blue or red (from neighbouring RBD) disks. (B), Sequence conservation the RBD determined by CONSERV showing strong conservation of residues forming the lipid binding pocket. (C) Surface representation of the RBD domains of the locked Spike with ACE2, CR3022 and S309 binding epitopes coloured in magenta, blue and cyan, respectively.

The locked conformation was observed alongside Spike trimers with “all-down” and “one-up” conformations in our single particle cryoEM data set. Inspection of the RBD density for the ‘down’ and ‘up’ conformations in our analyses and those reported previously (Huo et al., 2020; Walls et al., 2020; Wrapp et al., 2020) showed no evidence of pocket binder. However, since the density for RBDs in the ‘up’ conformation was of poor quality, we performed focused refinement of our data, isolating ‘up’ RBDs and their associated adjacent NTDs. This yielded an improved map with a resolution of 3.7 Å for these domains, sufficient to unambiguously assign side chains (Figure S3). The characteristic structure of residues D364-F374 confirmed no fatty acid was bound. Thus, the binding pocket is only found in the ‘locked’ conformation.

### Hydrophobic molecules perturb stability of the prefusion Spike

We hypothesized that hydrophobic molecules might replace the bound pocket factor (or occupy empty pockets). If the pocket factor modulates stability of the ‘locked’ trimer these events might alter the stability of the prefusion Spike. We therefore assembled a range of potential ligands which were tested in a thermofluor assay (Walter et al., 2012) (Figure S5, Methods). A number of molecules were assayed, based on the observed density in the pocket, being mainly fatty acids, pocket factors designed to replace fatty acids in enterovirus capsids (Tuthill et al., 2010) and compounds observed to bind Notum, which harbours a lipid binding pocket reminiscent of the one reported here (Kakugawa et al., 2015). The Spike exhibited two distinct melts, the first at ~53 °C, presumably a structural rearrangement, and a second melt, presumably the total denaturation of the protein, at ~68 °C. A number of compounds significantly reduced the first melting temperature (by 1.0 °C or more relative to a Spike with DMSO control). For both saturated and unsaturated fatty acids there was a clear trend with chain length (Figure S5). However no compounds showed a significant stabilisation of the Spike and we have no evidence that the compounds bind in the RBD pocket.

## Discussion

### Biological relevance of the ‘locked’ conformation

We have described a well-ordered closed conformation of prefusion Spike that appears to result from the incorporation of an endogenous ‘pocket-factor’. This ‘locked’ Spike conformation is stabilized by a series of specific, conserved interactions which tightly pack the pocket-filled RBD and NTD around the molecular 3-fold axis. It appears that RBD interactions generated in this structure stabilize regions of S2 involved in the structural transition to the postfusion state, notably the turn at residues 986-987 and the region adjacent to the fusion peptide (827-855), thereby preventing premature conversion of the prefusion state. These data, together with the recent observation of a similar structure in wild-type Spike ectodomain (Cai et al., 2020), strongly suggest that before recognition by a cellular receptor, the ‘locked’ conformation will be the predominant prefusion form on the viral surface. We believe that this has escaped notice primarily because attention has focused on a so-called stabilized Spike ectodomain construct, that has been engineered to prevent the switch to helical conformation in residues 986 and 987. Unwittingly these changes actually destabilize the ‘locked’ Spike, by abrogating the attachment of the region on the RBD which in the prefusion ‘all-down conformation’ appears to interact with the turn between the HR1 and CH helices forming a ‘cap’ or ‘lid’ to prevent the jack-knife helix extension and conversion to the postfusion form. Whilst the chemical identity of the lipid molecule we see bound in the pocket remains speculative, C18 fatty acids fit well with unsaturated Omega-3 or Omega-6 likely. It should also be borne in mind that the fatty acids available in the expression system might dictate what is recruited into the pocket, as has been observed for the analogous case of picornaviruses when VLPs of mammalian viruses were produced in insect or plant cells (Marsian et al., 2017; Ren et al., 2015), but note that our expression system uses mammalian cells. Fatty acid interactions between adjacent RBDs seem to play a key role in the stability of the ‘locked’ conformation. Interestingly fatty acids appear to bind to conserved regions of the Spike of an α-coronavirus (Kirchdoerfer et al., 2020), although binding does not occur in a pocket, and an array of hydrophobic compounds have been identified as stabilizers for MERS-CoV (ß subgroup C) (Su et al., 2016). The conservation of the residues forming the pocket we have identified (Figure 3) suggests that this locked conformation may occur across the β-coronaviruses as a ‘resting conformation’. This seclusion of the RBD might facilitate release of the virus from the cell and there would also be a selective advantage in keeping the immunogenic sites, including the ACE2 binding site, as inaccessible as possible. It is possible that the lipid then slowly diffuses out or a trigger, for instance engagement of an abundant attachment receptor (such as sialic acid or heparan sulphate), might facilitate the release of the fatty acid, initiating the switch to the ‘open’ state and presentation of the ACE2 binding site. Our observations also prompt the suspicion that production and purification protocols currently used may remove most of the lipid, contributing to the observed instability of the Spike.

The “locked” Spike conformation has considerable potential relevance for therapeutic design. A mechanism, whereby pocket-bound endogenous fatty acids stabilize a metastable state ahead of cell entry is analogous to the strategy used by some other RNA viruses, namely enteroviruses, where release of pocket-bound fatty-acids initiates a conformational change which primes the virion for genome release (Wang et al., 2012), and pocket factor mimics that fail to release have potential as drugs (De Colibus et al., 2014). In the case of Spike, we suggest that pocket binders which either stabilize or destabilize the Spike would offer a plausible route to a therapeutic, and we demonstrate that several compounds significantly destabilize Spike. The pocket is therefore a putative target for antiviral design. Knowledge of the mechanism of stabilization of the prefusion form of Spike is also important for understanding the context of epitopes presented *in vivo* and hence optimisation of presentation in non-classical vaccines. It is remarkable that the locked state of Spike serves to hide three key neutralizing epitopes, namely those of (i) ACE2 competing antibodies, for instance (Wu et al., 2020), (ii) antibodies that bind the S2 ‘capping’ epitope (exemplified by CR3022 (Huo et al., 2020)), and (iii) the extremely potent S309 antibody epitope (Pinto et al., 2020) (the major change in the conformation of the N343 glycan would be expected to disrupt antibody binding as the sugar contributes part of the epitope, Figure 2D, 3C). This suggests that a vaccine which presents these epitopes effectively, such as one using polyvalent displayed RBD domains might be more effective than a genuinely stable (‘locked’) prefusion Spike trimer. In summary, our findings are a further step towards understanding the structural dynamics of this complex and rather unstable molecule with implications for vaccines and antiviral therapy.

## METHODS

No statistical methods were used to predetermine sample size. The experiments were not randomized, and investigators were not blinded to allocation during experiments and outcome assessment.

### Genetic constructs of spike ectodomain

The gene encoding amino acids 1-1208 of the SARS-CoV-2 spike glycoprotein ectodomain, with mutations of RRAR > GSAS at residues 682-685 (the furin cleavage site) and KV > PP at residues 986-987, as well as inclusion of a T4 fibritin trimerisation domain, a HRV 3C cleavage site, a His-8 tag and a Twin-Strep-tag at the C-terminus. As reported by Wrapp *et al*. (Wrapp et al., 2020). All vectors were sequenced to confirm clones were correct.

### Protein production

Recombinant Spike ectodomain was expressed by transient transfection in HEK293S GnTI-cells (ATCC CRL-3022) for 9 days at 30 °C. Conditioned media was dialysed against 2x phosphate buffered saline pH 7.4 buffer. The Spike ectodomain was purified by immobilized metal affinity chromatography using Talon resin (Takara Bio) charged with cobalt followed by size exclusion chromatography using HiLoad 16/60 Superdex 200 column in 150 mM NaCl, 10 mM HEPES pH 8.0, 0.02% NaN_3_ at 4 °C.

### Complex Preparation and Cryo-EM Data Collection

Purified spike protein was buffer exchanged into 2 mM Tris pH 8.0, 200 mM NaCl, 0.02 % NaN_3_ buffer using a desalting column (Zeba, Thermo Fisher). A final concentration of 0.2 mg/mL was incubated with CR3022 Fab (in the same buffer) in a 6:1 molar ratio (Fab to trimeric spike) at room temperature. Aliquots were taken at 50 minutes. Immediately an aliquot was taken 3 μL of it was applied to a holey carbon-coated 200 mesh copper grid (C-Flat, CF-2/1, Protochips) that had been freshly glow discharged on high intensity for 20 s (Plasma Cleaner PDC-002-CE, Harrick Plasma) and excess liquid removed by blotting for 6 s with a blotting force of −1 using vitrobot filter paper (grade 595, Ted Pella Inc.) at 4.5 ºC, 100 % relative humidity. Blotted grids were then immediately plunge frozen using a Vitrobot Mark IV (Thermo Fisher). The analysis of the CR3022 bound fractions were analysed and published in an independent paper (BioRxiv: https://doi.org/10.1101/2020.05.05.079202).

### Cryo-EM Data collection and processing

Data were collected on a Titan Krios G2 (Thermo Fischer) at 300 kV). Automated data collection was setup in EPU 2.5 using a K3 (Gatan) direct electron detector operating in super-resolution mode at 0.415 Å per pixel and a GIF Quantum energy filter (Gatan) with a 20 eV slit. A defocus range of −0.8 μm to −2.6 μm and total dose of ~ 42.0 e^-^/Å^2^ across 40 frames was applied to the sample. Sample-specific data collection parameters are summarized in Table S1.

Motion correction of super-resolution raw movies were processed in Relion 3.1 (Zivanov et al., 2018), using a five-by-five patch-based alignment, keeping all the frames and a two time data binning, resulting in a final pixel size of 0.83 Å per pixel. The contrast transfer function (CTF) of full-dose, non-weighted micrographs was estimated using Gctf-v.1.06 (Zhang, 2016). Poor-quality images were discarded after manual inspection. Particles were blob picked in Cryosparc v.2.14.1 (Punjani et al., 2017) and after inspection of 2D classes, two sets of classes of interest were selected to generate templates for complete particle picking. A total number of 3,423,951 particles were picked and a final number of 468,112 particles were selected after two rounds of 2D classification. Three ab-initio models were generated and refined using heterogenous refinement. One good class containing around 328,000 particles was refined using non-uniform refinement to 3.3 Å and particles were exported to RELION 3.1 using pyEM csparc2star.py scripts (Asarnow et al., 2019). Bayesian polishing and per particle CTF refinement with beam tilt correction improved map resolution to 3.0 Å. For the 3D variability analysis, polished particles and model were imported into cryoSPARC v.2.14.0 and refined using the non-uniform refinement. The 3D variability analysis was then performed on the full map, asking for solving the three main modes on a structure low pass filtered to 6 Å. The results of this principal component analysis (PCA) were clustered in ten sub-populations and models were reconstructed for each of the individual clusters. One cluster was showing what we named the ‘locked’ conformation which was refined using the non-uniform refinement and C3 symmetry to 3.0 Å from 36,000 particles. In parallel, all the particles having a strong density for one of the RBD in the ‘up’ conformation were selected and refined using the non-uniform refinement and C1 symmetry to 3.1 Å from 191,000 particles. Local refinement after particle subtraction was performed on the RBD ‘up’ and some part of the S2 to allow good image alignment and another one focusing on the other part of the Spike, which were refined to 3.6 Å and 3.0 Å, respectively. Structures were modelled by first rigid body fitting the PDB ID, 6VYB (Walls et al., 2020) for the “up” conformation and by rigid body fitting individual domains for the “locked” conformation into the locally filtered map using UCSF Chimera (Pettersen et al., 2004) One cycle of rigid body real space refinement followed by manual adjustment in Coot (Emsley and Cowtan, 2004) was performed to correctly position the Cα chain into the density. Finally, cycles of PHENIX (Liebschner et al., 2019) real space refinement and manual building in Coot (Emsley and Cowtan, 2004) were used to improve model geometry. Map-to-model comparison in PHENIX (Liebschner et al., 2019) mtriage validated that no over-fitting was present in the structures. Model geometry was validated for all the refined models using MolProbity (Davis et al., 2004). All map and model statistics are detailed in Table S1.

### Thermofluor assays

were performed in triplicate in white 96-well PCR plates (4titude) using a RT-PCR machine (MX3005P, Agilent) with excitation and emission filters of 492 and 585nm. Experiments of 50 µL well volumes contained 0.2 µM protein (1-1.5 µg), a final dye concentration of 3X (SYPRO orange, Invitrogen) and 0.1M HEPES pH7.5, 0.1M NaCl. All screening compounds were dissolved in DMSO and used in large (50 00up to ~1000-fold) molar excess with a final concentration of 4% DMSO. A smaller excess of compound (~50-fold) was used if the background signal was seen to be so high that the melts could not be clearly observed: this was the case for the unsaturated fatty acids for example. To limit the self-quenching effect of the dye at elevated temperatures an “expanding sawtooth” profile was used in which the sample was held for 30 seconds at an incrementally increasing temperature from 25 to 98 °C before reading the fluorescence at 25 °C. Raw fluorescence data were exported and analysis was performed by fitting a 5-parameter sigmoid curve using the online software JTSA (Sacchi et al., 2015). The Tm was taken as being the midpoint of the curve. Further analysis was performed in Excel spreadsheets and Prism.

### Data availability

All data are available from the corresponding authors and/or included in the manuscript or Supplementary Information. Cryo-EM density maps with the corresponding atomic coordinates have been deposited in the Electron Microscopy Data Bank and the Protein Data Bank, respectively, with accession codes EMD-XXX, PDB ID XXX ‘locked’ Spike (C3), EMD-XXX, PDB ID XXX (‘locked’ Spike RBD/NTD focused), EMD-XXX, PDB ID XXX (‘all down’ Spike (C3)), EMD-XXX, PDB ID XXX (‘one up’ Spike (C1)), EMD-XXX, PDB ID XXX (‘one up’ Spike RBD/NTD focused), EMD-XXX, PDB ID XXX (‘one up’ Spike core/RBD down focused).

## MAIN TEXT STATEMENTS

## Acknowledgements

This work was supported by a grant from the CAMS-Oxford Institute to D.I.S. E.E.F and J. R. are supported by Wellcome (101122/Z/13/Z), Y.Z. by Cancer Research UK (C375/A17721) and D.I.S. and E.E.F. by the UK Medical Research Council (MR/N00065X/1). The Chinese Academy of Medical Sciences (CAMS) Innovation Fund for Medical Science (CIFMS), China (grant number: 2018-I2M-2-002) supports DIS. T.M. is supported by Cancer Research UK grants C20724/A14414 and C20724/A26752. R.R. was supported by a Wellcome PhD Training Programme 102 164/B/13/Z. This is a contribution from the UK Instruct-ERIC Centre. The Wellcome Centre for Human Genetics is supported by the Wellcome (grant 203141/Z/16/Z). The authors would like to thank Diamond Light source for access and support of the cryo-EM facilities at the UK national Electron Bio-Imaging Centre (eBIC), Proposal BI26983-2, both COVID-19 Rapid Access. Further EM provision was provided through the OPIC electron microscopy facility which was founded by a Wellcome JIF award (060208/Z/00/Z) and is supported by a Wellcome equipment grant (093305/Z/10/Z). Computation used the Oxford Biomedical Research Computing (BMRC) facility, a joint development between the Wellcome Centre for Human Genetics and the Big Data Institute supported by Health Data Research UK and the NIHR Oxford Biomedical Research Centre. Part of this work was supported by Wellcome administrative support grant (203141/Z/16/Z). We thank Yvonne Jones for advice and comments on the manuscript. Huge thanks to the teams, especially at the Diamond Light Source and Division of Structural Biology, Oxford University that have enabled work to continue during the pandemic.

## Author Information

These authors contributed equally: L. Carrique, H.M.E. Duyvesteyn, T. Malinauskas, Yuguang Zhao.

## Contributions

T.M., Y.Z., D.Z. and J.H. prepared the samples. H.M.E.D., R.R.R and P.N.M.S. performed cryo-EM sample preparation. J.Radecke performed cryo-EM data collection. H.M.E.D. and L.C. processed cryo-EM data. T.S.W. performed thermofluor expriments and analysed the data. L.C., H.M.E.D., J.R., T.S.W, E.E.F. and D.I.S. analysed the results and wrote the manuscript. All authors read and approved the manuscript.

**Correspondence and requests for materials** should be addressed to DIS/EEF

