## Supplementray information for "The SARS-CoV-2 Spike harbours a lipid binding pocket which modulates stability of the prefusion trimer"

### Supplementary information

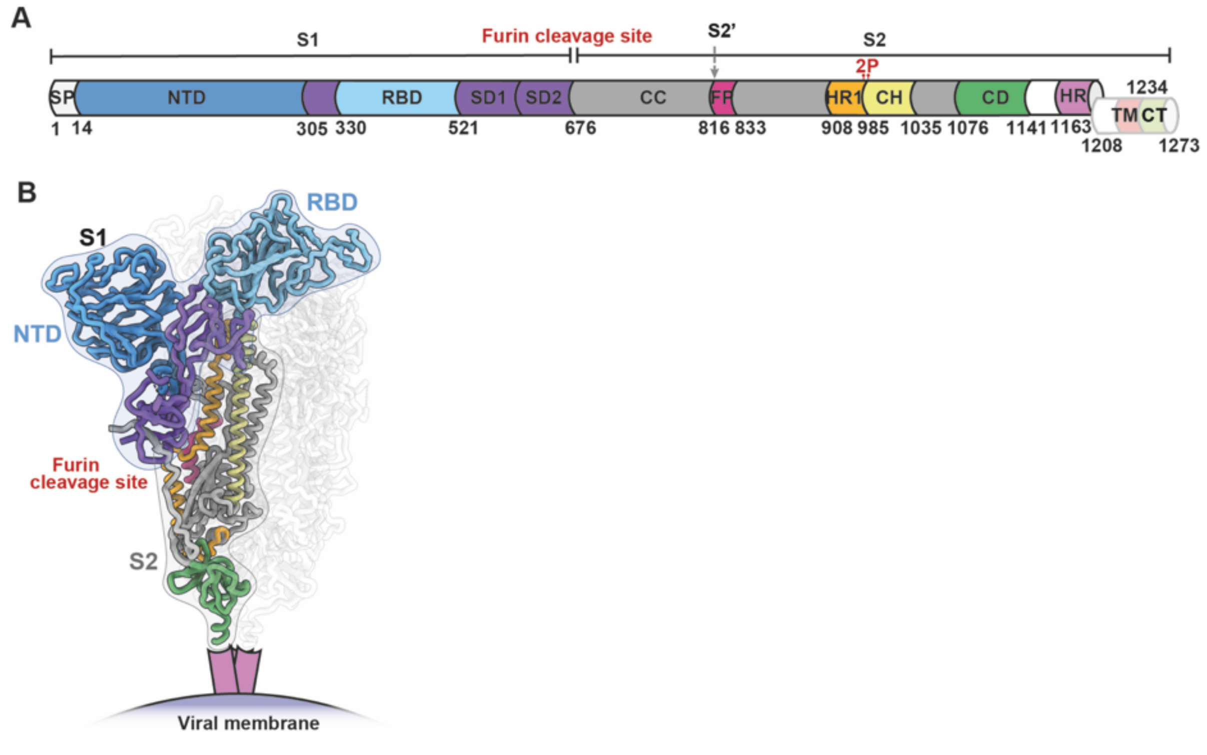


**Figure S1. Structural organisation of the SARS-CoV-2 spike glycoprotein.** (A) Domain mapping of the Spike, SP, signal peptide; RBD, receptor binding domain; SD1 and SD2, subdomains 1 and 2; CC, central Core; FP, fusion peptide; HR1, heptad repeat 1; CH, central helix; CD, connector domain; HR2, heptad repeat 2; TM, transmembrane domain; CT, cytoplasmic tail. (B) Side view of the 'locked' spike trimer with one protomer brightly coloured by domains.

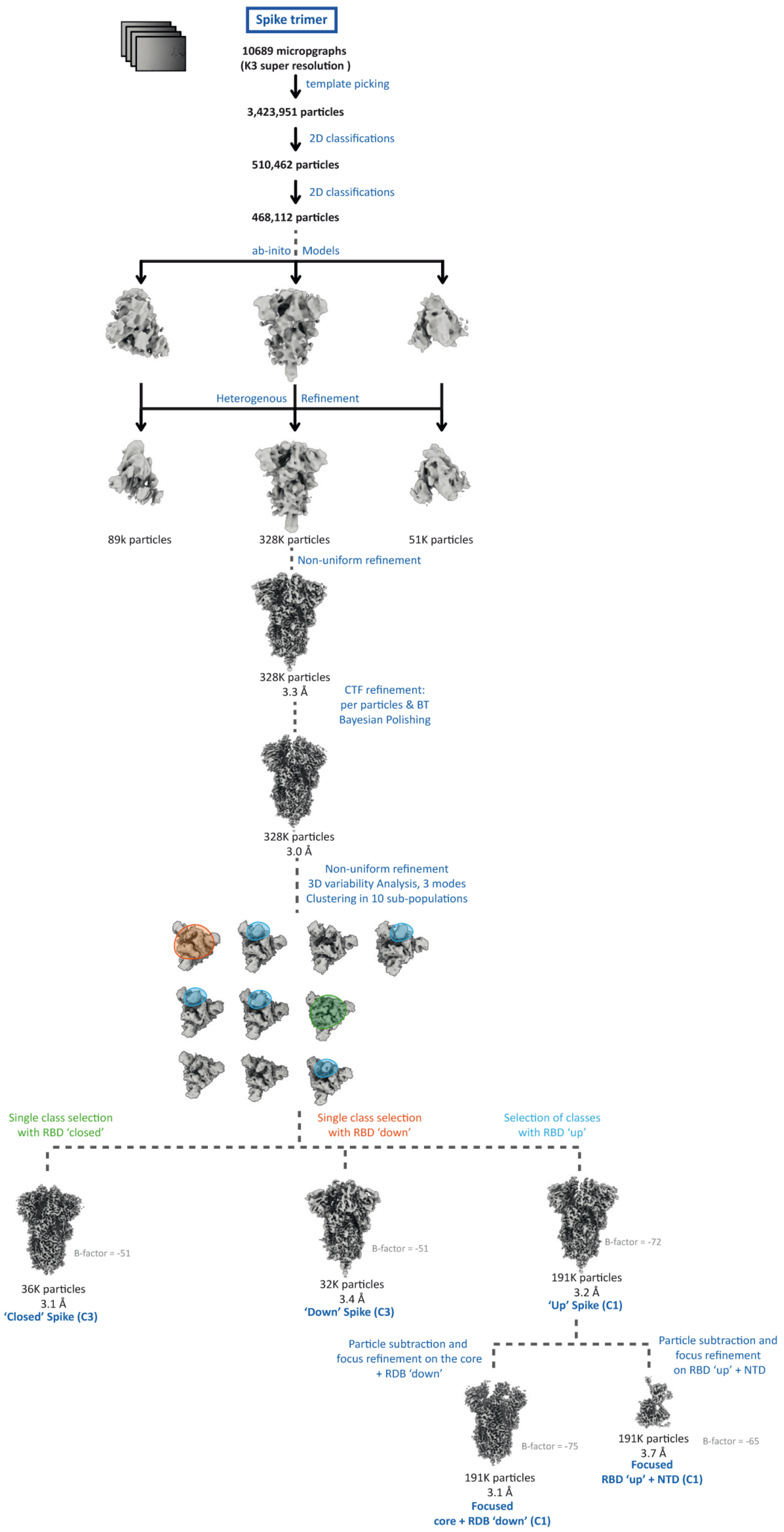


**Figure S2. Data collection, processing and analysis scheme.** Flowchart for the processing and classification of the trimeric Spike.

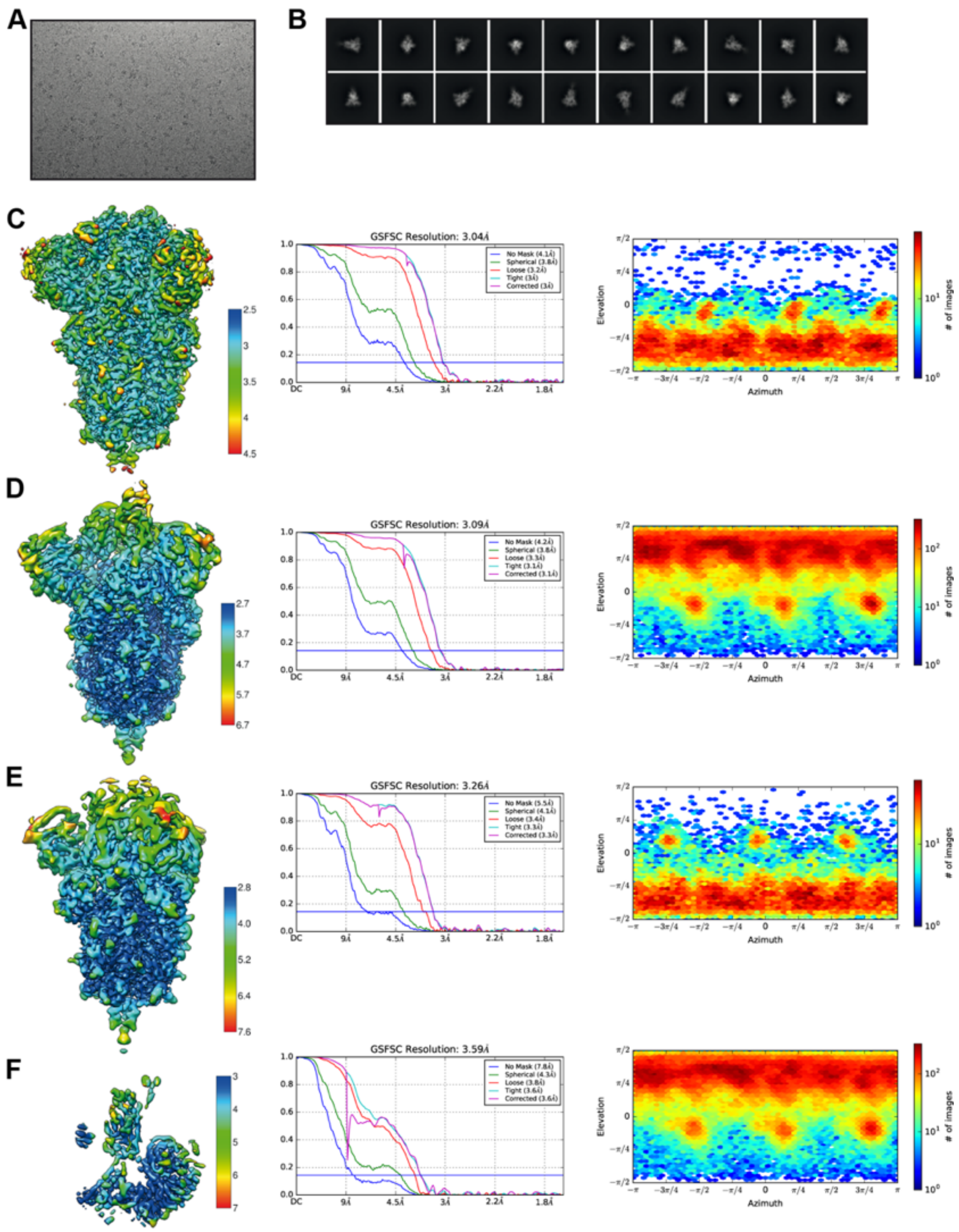


**Figure S3. Single-particle cryo-EM analysis.** (A) Representative micrograph of SARS-CoV-2 spike embedded in vitreous ice. (B) Representative 2D class averages of SARS-CoV-2 spike. (C-F) Data analysis for SARS-CoV-2 spike ‘Locked’ (C) ‘up’ (D) ‘down (E) and RBD-up focused (F) reconstructions. 3D reconstruction locally filtered and coloured according to local resolution (left panel). FSC curve indicating overall map resolution (middle panel). Angular distribution of particles used in the final reconstruction (right panel).

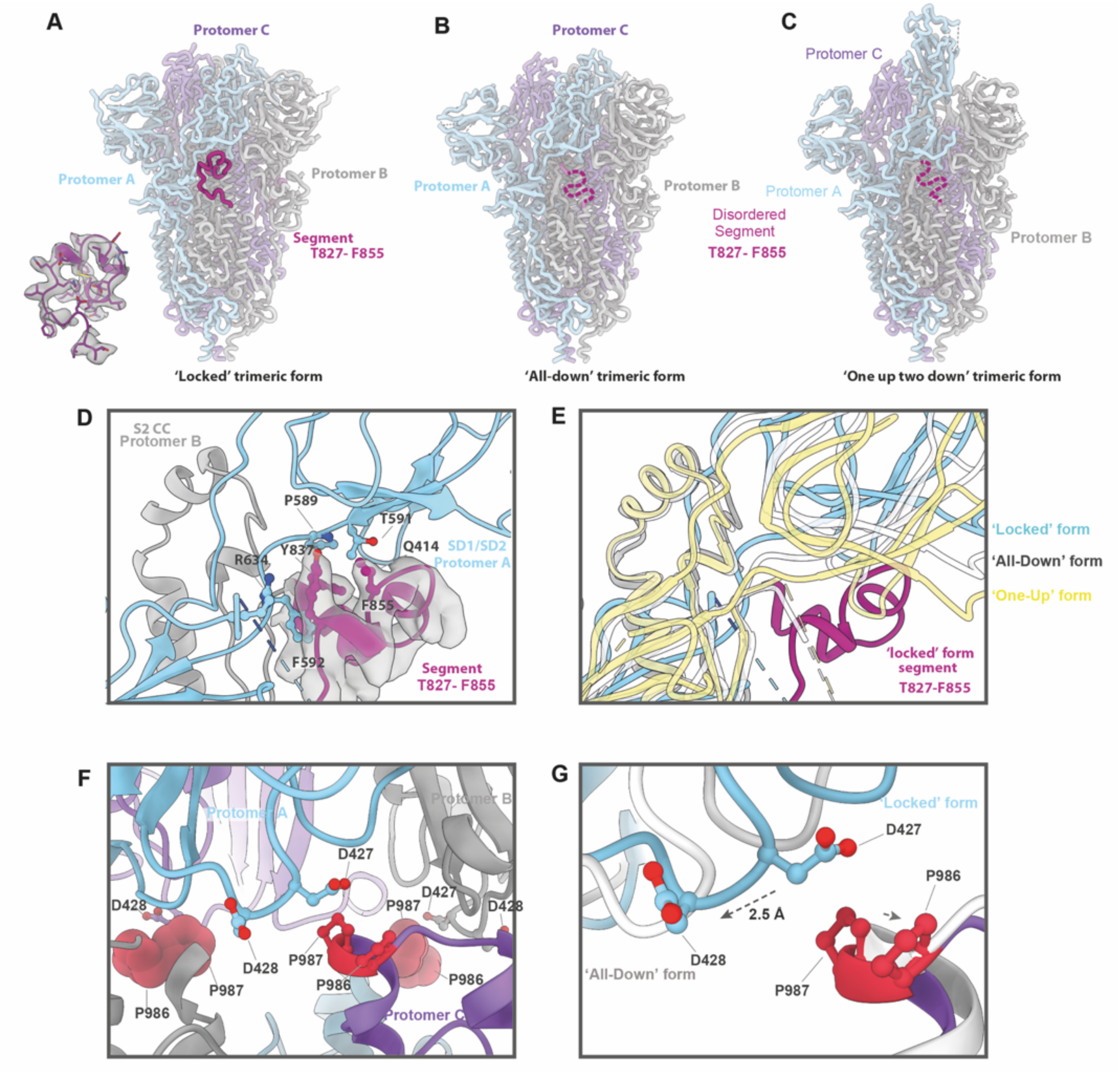


**Figure S4. Segment T827-F855 that mediates contact between S1 and S2 is disordered in the 'all down' and 'one up' states.** (A-C), Side views of the 'locked' (A), the 'all down' (B) (PDB ID, 6VXX) and the 'one up two down' (C) trimeric forms. The insertion at left bottom corner of panel (A) showing the density of T827-F855 in the 'locked' form. Segment T827-F855 is coloured in purple and highlighted with a dashed line in the 'all down' and 'one up'

forms where it is disordered. (D) Closeup of the interactions between T827-F855 segment of protomer B (grey) and SD1/SD2 of protomer A (cyan). (E) Overlay of the ‘all down’ (white) and ‘one up’ (yellow) forms showing the movement of SD1 in these two forms compared to the ‘locked’ form (purple). (F) Close view of K986P and V987P mutations (red) within the ‘locked’ trimeric form highlighting their proximity with D427 and D428. (G) Movement of the SD1 domain between the ‘locked’ form (cyan and purple) and the ‘all down’ form (grey) relative to K986P and V987P mutation sites (red).

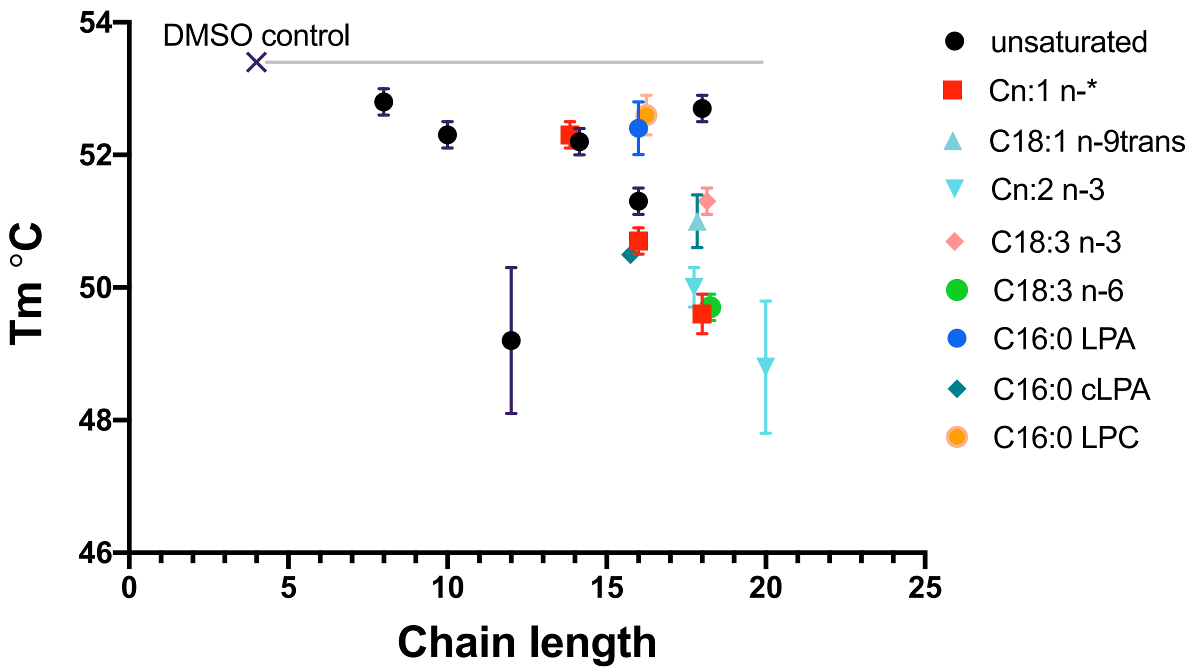


**Figure S5. The effect on Spike stability of fatty acids of different chain lengths**.

The first melting temperature is shown (see methods). The control uses Spike with the same concentration of DMSO as used as solvent for the lipids, but contains no lipids. Error bars are 1 standard deviation.

| | 'Locked' Spike (C3) | 'ALL Down' Spike (C3) | 'One Up' Spike (C1) | RBD 'One Up' focused (C1) |
|---|---|---|---|---|
| **Data collection** | | | | |
| Microscope | Titan Krios (eBIC) | | | |
| Voltage (kV) | 300 | | | |
| Detector | Gatan K3 | | | |
| Recording mode | Super Resolution | | | |
| Magnification | 130,000 | | | |
| Movie/micrograph pixel size (Å) | 0.415 | | | |
| Dose rate (e-/A$^2$/sec) | 20.7 | | | |
| Number of frames per movie | 40 | | | |
| Movie exposure time (s) | 2 | | | |
| Total dose (e-/Å$^2$) | 42 | | | |
| Defocus range (um) | -0.8 to -2.6 | | | |
| Volta Phase Plate | no | | | |
| **EM data processing** | | | | |
| Number of movies/micrographs | 10941 | | | |
| Box size (px) | 540 | | | |
| Particle number (total) | 3,423,951 | | | |
| Particle number (post 2D) | 468,112 | | | |
| Particle number (post 3D) | 328,000 | | | |
| Particle number (used in final map) | 36K | 32K | 191K | 191K |
| Symmetry | C3 | C3 | C1 | C1 |
| Map resolution (FSC 0.143) | 3.1 | 3.3 | 3.2 | 3.7 |
| Map sharpening B-factor (A$^2$) | -51 | -59 | -72 | -65 |
| Model Building and Validation | | | | |
| Initial model used | 6VXX | / | 6VYB | 6VYB |
| **Model composition** | | | | |
| Non-H protein atoms | 52026 | / | 43559 | 7552 |
| Protein residues | 3267 | / | 2875 | 466 |
| Ligands STE | 3 | / | / | / |
| Ligands NAG | 66 | / | 59 | 7 |
| **RMSD from ideal** | | | | |
| Bond length (Å) | 0.01 | / | 0.02 | 0.01 |
| Bond angles (°) | 0.72 | / | 1.20 | 0.92 |
| **Validation** | | | | |
| Molprobity score | 1.7 | / | 1.5 | 2.2 |
| Clashscore | 4.8 | / | 6.4 | 11.9 |
| Rotamers outliers (%) | 0.4 | / | 0.7 | 0.2 |
| FSC (0.5) model-vs-map | 3.1 | / | 3.2 | 3.8 |
| CC model-vs-map (masked) | 0.87 | / | 0.8 | 0.78 |
| **Ramachandran plot** | | | | |
| Favored (%) | 94 | / | 97 | 86 |
| Allowed (%) | 6 | / | 2 | 14 |
| Outliers (%) | 0 | / | 0 | 0 |

**Table S1 Cryo-EM data collection, refinement, and validation statistics**

**Video S1 Morph between the 'locked', the 'down' and the 'up' conformation of the SARS-CoV-2 prefusion spike.** The movie shows the structural rearrangement occurring between the 'locked' and the 'down' conformation and the between the 'down' and the 'up' conformation.

**Video S2 Morph between the RBD and the RBD bound the palmitoleic acid.** The movie shows the structural rearrangement occurring between the RBD and the RBD bound the palmitoleic acid with segment$_{D364-F374}$ highlighted in orange.
